# BzAM terpolymer series with a step-wise systematic change in hydrophobicity, provides a nuanced platform for functional analysis of GPCRs in native-like nanodiscs

**DOI:** 10.64898/2026.05.20.726474

**Authors:** Gestél C. Kuyler, Rebecca J. Murray, Farhaan Napier Khwaja, Joseph Gunner, Bert Klumperman, David Poyner, Hoor Ayub, Mark Wheatley

## Abstract

Detergent-free extraction of membrane proteins using polymers directly into nanodiscs from the cell membrane has been used widely in recent years. Since the first use of poly(styrene-*co*-maleic acid) (SMA), numerous related polymers have been developed that differ in chemical architecture and nanodisc characteristics, each capable of influencing the structural and functional properties of the encapsulated membrane protein and its surrounding lipids. Identifying an optimal solubilising polymer, therefore, requires consideration not only of extraction efficiency but also compatibility with downstream applications and analyses. Polymer series in which a single parameter is systematically varied provide a valuable, nuanced tool for optimising nanodisc utility in downstream applications. This study utilises a chemically defined series of poly(styrene-*co*-maleic acid-*co*-(*N*-benzyl)maleimide) (BzAM) terpolymers that exhibit a stepwise, systematic increase in hydrophobicity. Using the human calcitonin gene-related peptide (CGRP) receptor as an exemplar class B1 G-protein-coupled receptor (GPCR), the ability of each BzAM terpolymer to solubilise the receptor from mammalian cell membranes was assessed. All members of the series successfully solubilised CGRP receptor, with solubilisation efficiency correlating positively with increasing hydrophobicity. Importantly, the receptor retained its characteristic high-affinity ligand-binding capability when encapsulated within the BzAM nanodisc, demonstrating that functional integrity is preserved following BzAM-mediated extraction and purification. These findings establish the BzAM terpolymer series as a systematic, tuneable, well-defined tool for the detergent-free solubilisation and functional investigation of GPCRs, and other membrane proteins, in near-native lipid environments.

**HIGHLIGHTS:**

- Stepwise-tuned poly(styrene-*co*-maleic acid-*co*-(*N*-benzyl)maleimide) (BzAM) terpolymers provide a chemically defined, hydrophobicity-controlled platform for detergent-free membrane protein extraction.
- All BzAM variants effectively solubilise the human calcitonin gene-related peptide (CGRP) receptor, with extraction efficiency increasing in line with terpolymer hydrophobicity.
- CGRP receptor maintains high-affinity ligand binding in BzAM nanodiscs, demonstrating preservation of ligand-binding function after solubilisation.
- The BzAM series provides a novel platform for studying G-protein-coupled receptors and other membrane proteins in near-native lipid environments, with the potential to deliver mechanistic insights and support future drug-discovery efforts.

**GRAPHICAL ABSTRACT:** 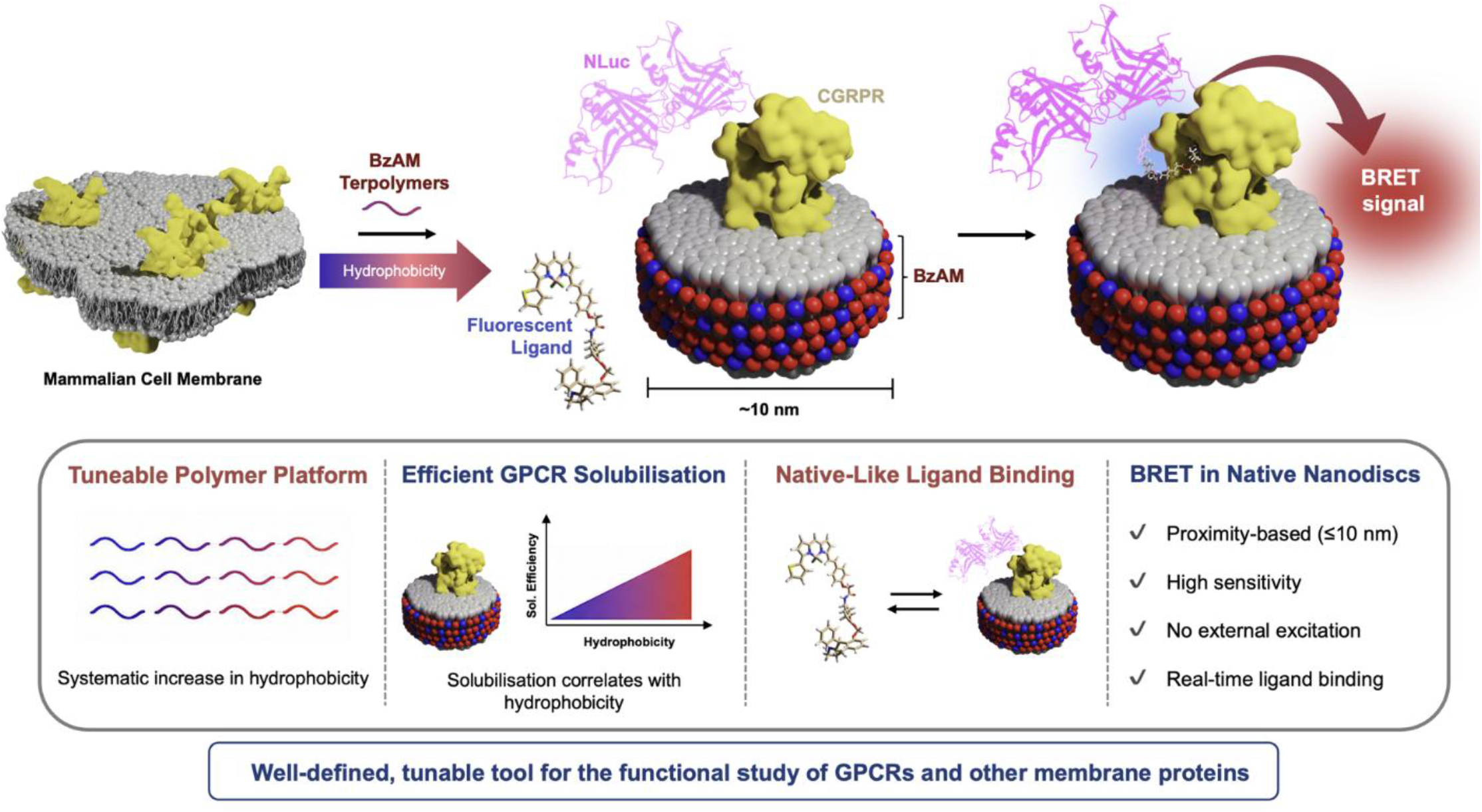

## 1. INTRODUCTION

The extraction of integral membrane proteins from the plasma membrane of cells while retaining their functional capability is challenging. Nevertheless, extraction and purification are routinely prerequisites to fully understanding the structure and associated molecular mechanisms of membrane proteins of interest. The largest family of membrane proteins in the human genome is the G-protein-coupled receptor (GPCR) family, with >800 members. GPCRs regulate a plethora of physiological processes and are the largest therapeutic target in humans, with *ca*. 35% of clinically-approved drugs targeting GPCRs (1). Therefore, as a family of membrane proteins, GPCRs are high-value targets for study.

Conventional detergent-based solubilisation effectively disrupts the lipid bilayer, extracting proteins into detergent micelles. However, by stripping away annular lipids, disrupting lipid–protein interactions and ablating membrane lateral pressure, this process is detrimental to membrane protein function and stability. Detergent-based solubilisation also removes the subtle lipid heterogeneity of the native bilayer, including lipid asymmetry between the bilayer leaflets (2,3); enrichment of certain lipids surrounding membrane proteins (4) and the existence of micro-domains with specific lipid enrichment, such as ‘lipid rafts’ (5). The term ‘functional paralipidome’ has been proposed to describe the preferred lipid microenvironments that solvate membrane proteins (6). Furthermore, for GPCRs, specific lipids have been reported to exert functional effects [reviewed in (7)], including regulation of ligand binding and intracellular signalling by cholesterol (8), allosteric regulation of active and inactive conformations by phosphatidylglycerol and phosphatidylethanolamine (9), stabilisation of active receptor conformation and enhanced G-protein selectivity by phosphatidylinositol 4,5 bisphosphate (10) plus modulation of GPCR conformational plasticity by phosphatidylserine (11) and anionic lipids (12). Clearly, the lipid context is important for membrane proteins in general, and for GPCR function in particular.

Polymer-stabilised nanodiscs, such as those created from poly(styrene-*co*-maleic acid) (SMA) copolymers (*i*.*e*., SMA lipid particles; SMALPs), offer a detergent-free alternative, retain the annular lipids (13–15) and facilitate downstream biophysical investigation (16–18). Despite significant advancements in recent years, the commonly used copolymer SMA2:1 (with a 2:1 styrene to maleic acid ratio) suffers from inherent drawbacks related to the synthetic strategy employed in its production. Conventional radical polymerisation results in broad molecular weight distributions, leading to a highly heterogeneous mixture of polymer chain lengths within a single sample and limited control over chain-end functional capabilities. This may present limitations in methodical evaluation and in gaining a molecular understanding of polymer-lipid interactions (19). Therefore, well-defined polymers are more reliable tools for investigating membrane proteins due to reduced uncertainties and improved uniformity within the system (20).

Controlled polymerisation techniques, such as reversible addition−fragmentation chain transfer (RAFT)-mediated polymerisation, allow rational polymer design with defined chemical characteristics and narrow molecular weight distributions. Recently, a novel series of poly(styrene-*co*-maleic acid-*co*-(*N*-benzyl)maleimide (BzAM) terpolymers was developed (21). These terpolymers were synthesised from the structurally defined, alternating base copolymer, poly(styrene-*alt*-maleic anhydride) (SMAnh). Partial modification of SMAnh with *N*-benzylamine produced a series of BzAM terpolymers, where increasing *N*-benzylmaleimide substitution systematically increased hydrophobicity throughout the series.

GPCRs are subdivided into families based on structural homology fingerprints (22). The receptor for the neuropeptide calcitonin gene-related peptide (CGRP), a class B1 GPCR, is a high-value target because it modulates vascular tone, inflammatory and metabolic responses, and is a major therapeutic target for the treatment of migraine (23). In this study, we investigate the ability of the individual polymers in the BzAM terpolymer series to extract the human CGRP receptor (CGRPR). We show that the CGRPR is effectively solubilised by the BzAM terpolymers, with the degree of extraction dictated by the hydrophobicity incorporated into the polymer architecture. Following solubilisation, representative nanodiscs generated with BzAM35 and containing Nanoluciferase (NLuc) tagged CGRPR were purified and assessed using a Nanoluciferase Bioluminescence Resonance Energy Transfer (NanoBRET) ligand binding assay (24,25), in which binding of a fluorescently labelled α-CGRP analogue to NLuc-CGRPR was quantified.

Bioluminescence Resonance Energy Transfer (BRET) is a sensitive method for monitoring receptor-ligand interactions in real time. In BRET assays, light emitted by a luciferase donor is transferred to a fluorescent acceptor when the two are within ∼10 nm, producing a measurable signal (26,27). This proximity requirement could make polymer-stabilised nanodiscs particularly well suited to BRET, as their nanoscale dimensions confine receptors and ligands within the distance range needed for efficient energy transfer. BRET offers high sensitivity with minimal photobleaching and outperforms Förster Resonance Energy Transfer (FRET)- and radioligand-based methods for studying GPCR binding dynamics. Although BRET-based assays are used for functional receptor studies in living cells and in copolymer encapsulated receptors, including class B1 GPCRs by Khwaja *et al*. for CGRP receptor and parathyroid hormone 1 receptor (28), this work represents BzAM-stabilised nanodiscs suitability for BRET application to quantify ligand binding. Given reported polymer-specific effects on GPCR conformation (29,30), verifying that BzAM-encapsulated receptors retain high-affinity ligand binding is essential. Here, we apply NanoBRET-based assay in BzAM polymer-stabilised nanodiscs to evaluate the function of (NLuc-tagged CGRPR using a fluorescently derivatised bespoke α-CGRP peptide, providing a controlled environment for probing GPCR ligand-binding ability in a native-like membrane-mimetic system. This proximity-based approach demonstrates that the ligand-binding capability of the CGRPR is retained when the receptor is encapsulated in a BzAM-stabilised nanodisc. This establishes the utility of the BzAM terpolymer systematic series as a valuable new tool for studying membrane proteins expressed by mammalian cells, particularly GPCRs.

## 2. MATERIALS AND METHODS

### 2.1. Materials

The copolymer SMA2:1 (M_w_ 7500 Da, *Ð* 2.50) was provided by Professor Timothy Dafforn (University of Birmingham, UK).

complete™ Mini EDTA-free protease inhibitor cocktail tablets (#11836170001, Roche), DMEM High glucose w/L-glutamine w/o sodium pyruvate (Biosera), Dulbecco’s modified Eagles medium/F-12 nutrient mixture (DMEM/F12) (#11330057, Thermofisher scientific), Geneticin (#10131027, Thermofisher scientific), IRDye® 800CW Donkey anti-mouse IgG Secondary antibody (#926-32212, LiCOR), Monoclonal Anti-HA antibody produced in mouse (#H9658-.2ML, Sigma Aldrich), Ni-NTA Agarose (#30210, Qiagen), Olcegepant (#4561, Tocris),

### 2.2. Fluorescent CGRP

A custom fluorescent α-CGRP analogue was constructed by substituting the native Asp3 of human α-CGRP with propargylglycine (Pra) to enable site-specific labelling following strategy previously reported for tetramethylrhodamine (TAMRA) labelling (31). The modified peptide was functionalised at Pra with BODIPY 630/650 via alkyne-azide cycloaddition, generating a BODIPY-CGRP ligand. Synthesis was performed using Biosynth custom peptide synthesis service. The peptide sequence and characterisation data were previously reported by Khwaja *et al*. (28).

### 2.3. CGRPR cDNA construct design

To achieve coordinated expression of CLR and RAMP1, bespoke bi-gene CGRPR constructs was synthesised in which both subunits are produced from a single transcript. A porcine teschovirus-1 2A (P2A) self-cleaving peptide sequence (ATNFSLLKQAGDVEENPGP) (32) was inserted between the human CLR and RAMP1 coding regions.

For NLuc-CGRPR construct, CLR was N-terminally tagged with NLuc to enable NanoBRET based ligand binding assays. The construct includes an N-terminal His_10_ tag and HA epitope, followed by the NLuc tag (NLuc-CGRPR only), the human CLR coding sequence, an AVI-tag (for CGRPR) or Twin-Strep tag (for NLuc-CGRPR), the P2A element, a FLAG tag, and the human RAMP1 coding region, terminating with a stop codon.

The custom cDNA was synthesised by GenScript (UK) and cloned into pcDNA 3.1(+) or pACMV-TetO vectors as described previously (33). Both CGRPR and NLuc-CGRPR construct designs and corresponding amino-acid sequences were as described by Khwaja *et al*. (28).

### 2.4. Cell culture

HEK 293S cells stably-expressing CGRPR or NLuc-CGRPR were routinely cultured in DMEM/F12 in the presence of G418 (0.5 mg·mL^-1^) supplemented with 10% (v/v) fetal bovine serum (FBS) in humidified 5% (v/v) CO_2_ in air at 37 °C. To induce expression of CGRPR, or Nluc-CGRPR, fresh DMEM/F12 without G418 and with 2 μg·mL^-1^ tetracycline was added for 48 h prior to harvesting.

### 2.5 Cell membrane preparation

Cells were harvested (900 × *g*, 5 min, 4 °C), resuspended in homogenisation buffer (10 mM Tris-base, 250 mM sucrose, 1 mM EDTA, pH 7.4) supplemented with protease inhibitor tablet and disrupted by 20 passes through a 21-gauge needle. After removal of cell debris (3,500 × *g*, 20 min, 4 °C), membranes were pelleted (100,000 × *g*, 1 h, 4 °C) and the wet pellet weight determined. Membrane pellets were resuspended in Membrane Suspension Buffer (MSB) (20 mM HEPES, 50 mM NaCl, 10% glycerol, pH 7.0) at 80 mg·mL^-1^ and aliquots stored at -80 °C.

### 2.6. Synthesis of the BzAM terpolymer series

RAFT-mediated polymerisation was used to synthesise the alternating base copolymer of poly(styrene-*alt-*maleic anhydride) (SMAnh 1:1). Basic hydrolysis of this yielded poly(styrene-*alt-*maleic acid) (SMA 1:1). Incremental modification of SMAnh 1:1 with varying molar amounts of *N*-benzylamine was followed by alkaline hydrolysis to produce the poly(styrene-*co*-maleic acid-*co*-(*N-*benzyl)maleimide) (BzAM) terpolymer series, as described by Kuyler *et al*. (21). The structure of BzAM polymers was confirmed by ^1^H NMR and ATR-FTIR spectroscopy. Quantitative ^13^C NMR spectroscopy was used to quantify the degree of benzylamine modification. Molecular weight and dispersity values (M_n_ 5000 Da; *Ð* 1.36) were obtained by size exclusion chromatography in DMF mobile phase calibrated with poly(methyl methacrylate) standards.

### 2.7. Solubilisation of cell membranes by polymers

Equal volumes of polymer stock solution (5.0% (w/v) in MSB) and cell membrane preparation (80 mg·mL^-1^) were mixed to yield a final polymer concentration of 2.5% (w/v) and a final membrane fraction concentration of 40 mg·mL^-1^. Following incubation for 2 h at 25 °C, samples were centrifuged (100,000 × *g*, 1 h, 4 °C; Sorvall WX Ultra 80 Series Ultracentrifuge) to separate solubilised receptors encapsulated in polymer-generated nanodiscs from insoluble material. Prior to SDS-PAGE analysis, insoluble fractions were resuspended to their original volume in MSB. Likewise, samples of membrane preparation were diluted appropriately and analysed alongside polymer-solubilised and insoluble fractions as a control to determine solubilisation efficiency.

### 2.8. SDS-PAGE and Western blot analysis

Samples were analysed by SDS-PAGE using pre-cast mini-protean TGX gradient (4-15%) gels (BioRad) run at 80 V for 1.5 h. Protein bands were visualised using SilverQuest Silver Staining Kit (Invitrogen) according to the manufacturer’s instructions.

For Western blot analysis, proteins were transferred from the SDS-PAGE gel to a nitrocellulose membrane, which after blocking with 5% skimmed milk in 0.1% PBST (phosphate-buffered saline with 0.1% v/v Tween20), was probed with primary anti-HA antibody overnight at 4 °C, followed by incubation with fluorescent secondary antibody for 1 h, protected from light. A LiCor Odyssey FC was used for visualisation, with densitometric analysis and image processing performed in Image Studio (version 5.2.5).

### 2.9. Receptor purification

Polymer-solubilised CGRPR-nanodiscs were mixed with Ni-NTA Agarose resin on a rotary shaker, overnight at 4 °C. The mixture was transferred to a gravity-flow column and the flow-through was collected. The resin was washed with 5 × 5 bed volumes (BV) of purification buffer (20 mM Tris, 200 mM NaCl, pH 8) supplemented with 20 mM imidazole, followed by 2 × 5 BV of purification buffer containing 40 mM imidazole, then 1 BV of purification buffer supplemented with 60 mM imidazole. CGRPR-containing BzAM lipid particles (BzAMLPs) were eluted from the resin with 6 × 0.5 BV of purification buffer supplemented with 300 mM imidazole. Elution fractions were pooled.

### 2.10. NanoBRET Ligand binding

CGRPR-BzAMLP nanodiscs in the pooled 300 mM imidazole elution fraction from the Ni-NTA column were washed with 3 × 20 mL BRET buffer (20 mM HEPES, 150 mM NaCl, 1mM EGTA, pH 7.4) in a spin-concentrator (Vivaspin 10 kDa MW cut-off, Sartorius, 1,800 × *g*) to remove imidazole from the sample, then concentrated to ∼500 μL. Ligand-binding assays were performed in white 384-well optiplates (PerkinElmer). NLuc-CGRPR-BzAMLPs were incubated with varying concentrations of BODIPY-CGRP, as indicated, for 45 min at 25 °C. Furimazine (10 µM) was then added and luminescence measured after 5 min using a CLARIOstar plate reader (BMG Labtech) with excitation at 460 nm (80 nm bandpass) and emission at 660 nm (100 nm bandpass). The raw BRET ratio was calculated as the ratio of long-wavelength to short-wavelength emission. Non-specific binding was determined in the presence of olcegepant (10 nM). Specific binding was then calculated by subtracting the raw BRET ratio of the olcegepant-treated sample (non-specific binding) from the raw BRET ratio of the vehicle-treated sample (total binding). Binding data were analysed using GraphPad Prism 8.0 (GraphPad Software Inc., San Diego, CA) and fitted with one-site nonlinear regression for specific binding.

## 3. RESULTS AND DISCUSSION

### 3.1. Effect of Polymer Amphiphilicity

The BzAM terpolymer series was designed to retain key characteristics of the widely used SMA2:1 copolymer, particularly with respect to its anionic nature and aromatic hydrophobic moieties, while overcoming limitations associated with its molecular weight heterogeneity. The goal was to develop well-defined polymers with effective solubilisation capability while offering enhanced control over molecular weight, molecular weight distribution, chemical structure, and chain-end functionalisation. Using RAFT-mediated polymerisation, the alternating copolymer SMAnh1:1 was synthesised as a precursor and partially derivatised with *N*-benzylamine to yield the BzAM series (Figure 1). Incrementally increasing the molar fraction of *N*- benzylmaleimide generated a series of structurally-defined terpolymers (BzAM05– BzAM40), in which substitution increased in 5% increments from 5% (BzAM05) to 40% (BzAM40), resulting in a systematic, stepwise increase in hydrophobicity across the series.

**Figure 1.**
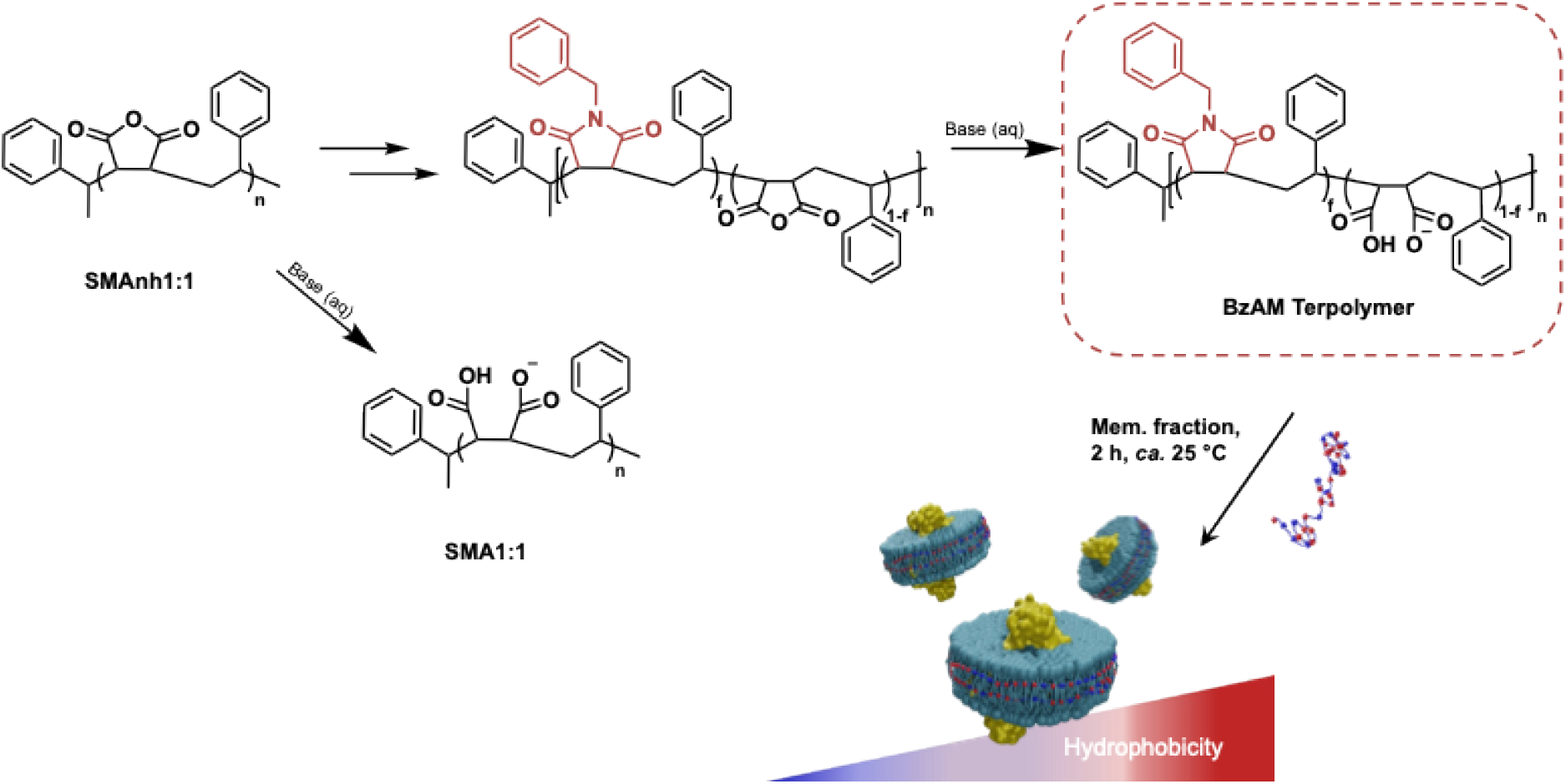
Synthetic overview of BzAM terpolymers from the alternating base copolymer, SMAnh. SMAnh, poly(styrene-alt-maleic anhydride; SMA, poly(styrene-co-maleic acid; BzAM, poly(styrene-co-maleic acid-co-(N-benzyl)maleimide.

The CGRPR is a heterodimer composed of a seven transmembrane-helix (TM) domain calcitonin receptor-like receptor (CLR), in complex with receptor activity-modifying protein-1 (RAMP-1), which has a single TM. A HEK 293 cell line was generated stably expressing the human CGRPR in which the CLR component incorporated an *N*-terminal (His)_10_ followed by an HA epitope-tag, to facilitate purification and downstream analysis, respectively, followed by nanoluciferase (NLuc). The ability of the individual terpolymers of the BzAM series to extract CGRPR from the cell membrane was investigated. Incubation of CGRPR-expressing cell membranes with the BzAM terpolymers at 2.5% (w/v) with mild agitation for 2 h at *ca*. 25 °C, resulted in the samples becoming clear, consistent with membrane solubilisation. Following ultracentrifugation (100,000 × *g*, 1 h) to separate the solubilised sample (supernatant) from the insoluble material (pellet), the ability of individual BzAM terpolymers to solubilise CGRPR was assessed by Western blot analysis, utilising the HA-tag on CLR. A characteristic band corresponding to CLR, migrating at 75 kDa, was observed across the BzAM terpolymer series (Figure *2*). Improvement in solubilisation efficiency resulted in an increase in band intensity in the solubilised fraction (Figure 2A) and a decrease in the band intensity in the insoluble fraction (Figure 2B). The base copolymer SMA1:1 exhibited low solubilisation efficiency (12 ± 1.9% mean ± s.e.m., n = 3), being approximately 8-fold lower than SMA2:1, consistent with previous observations (21,34).

**Figure 2.**
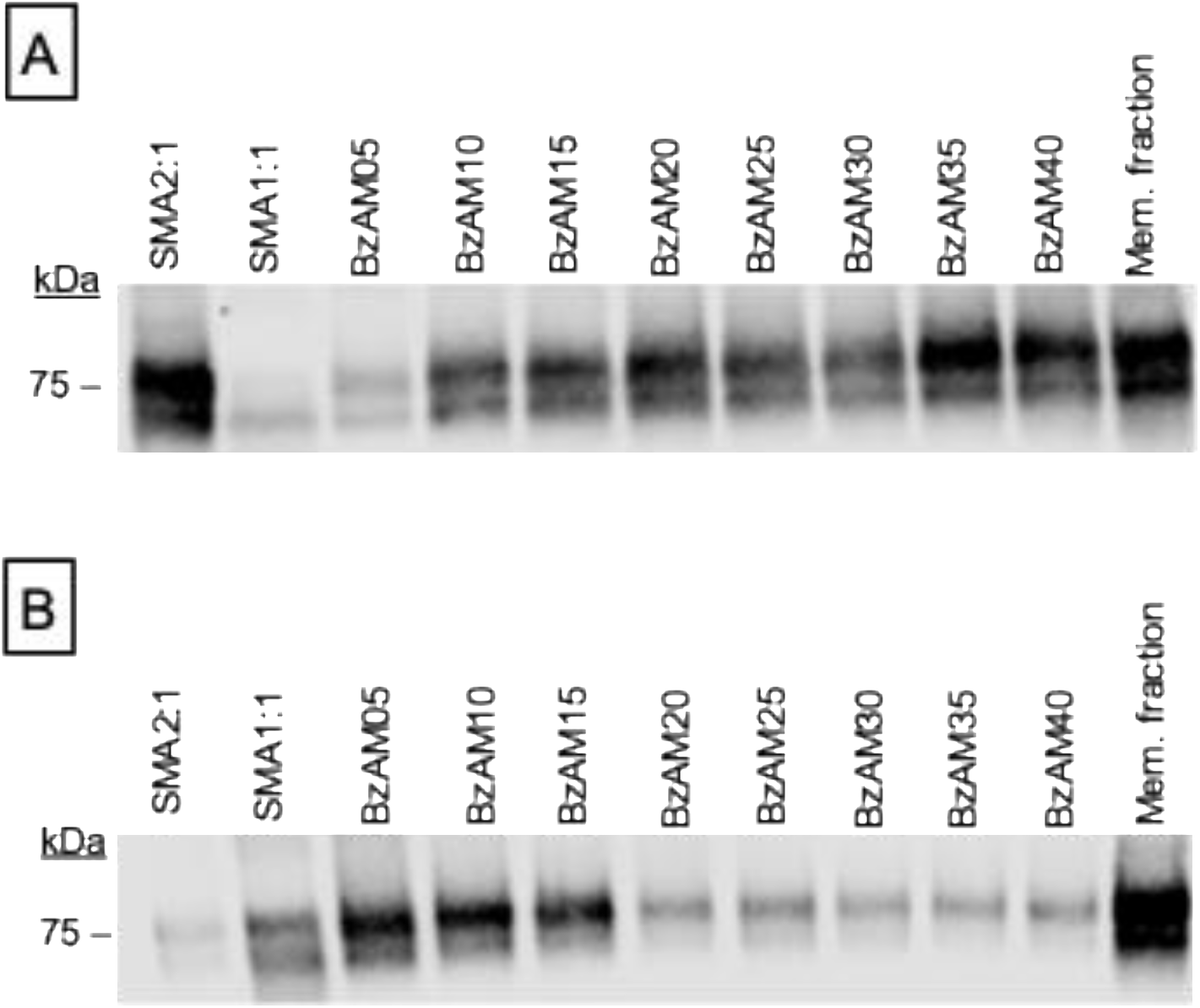
Solubilisation of CGRPR by polymers. Following incubation of CGRPR-expressing cell membranes with the stated polymers (2.5% (w/v) polymer, 2 h, ca. 25 °C), samples were centrifuged (100,000 × g, 1 h, 4 °C) to separate solubilised receptor (supernatant) from insoluble material (pellet). Western blot analysis of the supernatant (panel A) and pellet (panel B) for each polymer is presented. The original membrane preparation (Mem. fraction) is shown for reference.

Densiometric analysis of the 75 kDa band in each of the BzAM-solubilised fractions was compared to the corresponding band in the original membrane preparation to determine the percentage of CGRPR solubilised by each terpolymer (Figure *3*).

Conversion of approximately 5% of the hydrophilic repeat units of SMA1:1 to *N-*benzyl maleimide (BzAM05) resulted in a significant increase in CGRPR solubilisation to 25 ± 2.5% (mean ± s.e.m. n = 3; Figure 3). Solubilisation efficiency increased to 61 ± 8.9% (mean ± s.e.m., n = 3) with BzAM10 and continued to increase throughout the series, up to BzAM40.

**Figure 3.**
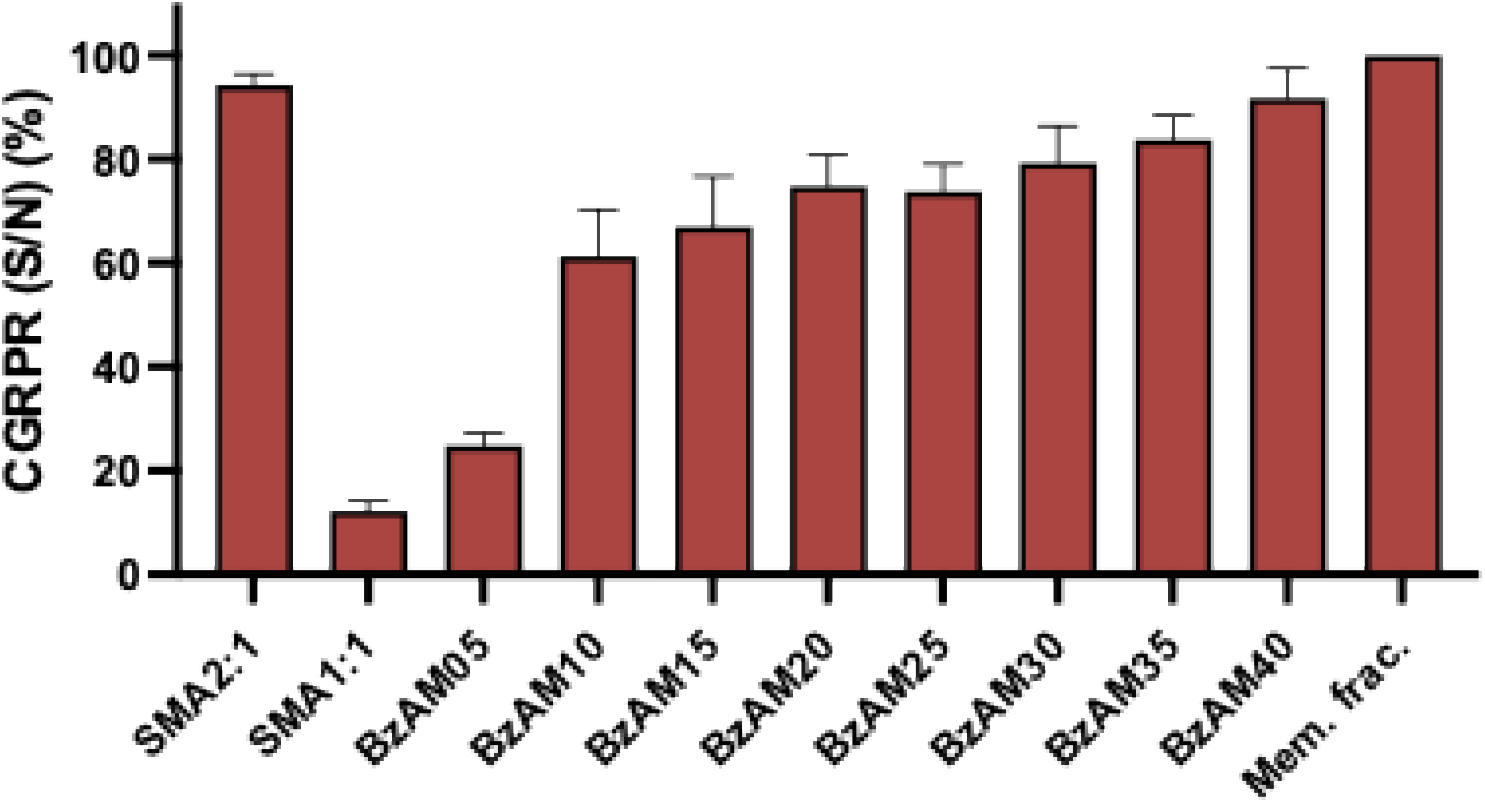
CGRPR solubilisation efficiency of polymers. For the polymers stated, the solubilisation efficiency was calculated as the percentage of CGRPR in the original membrane preparation (Mem. frac.) detected in the soluble fraction (S/N), determined by densitometric analysis of western blots. Data are mean ± s.e.m. (n = 3).

It was noteworthy that for the terpolymers in the series between BzAM10 and BzAM40 inclusive, there was a clear correlation between the CGRPR solubilisation efficiency and the hydrophobicity of the terpolymer (Figure 4).

**Figure 4.**
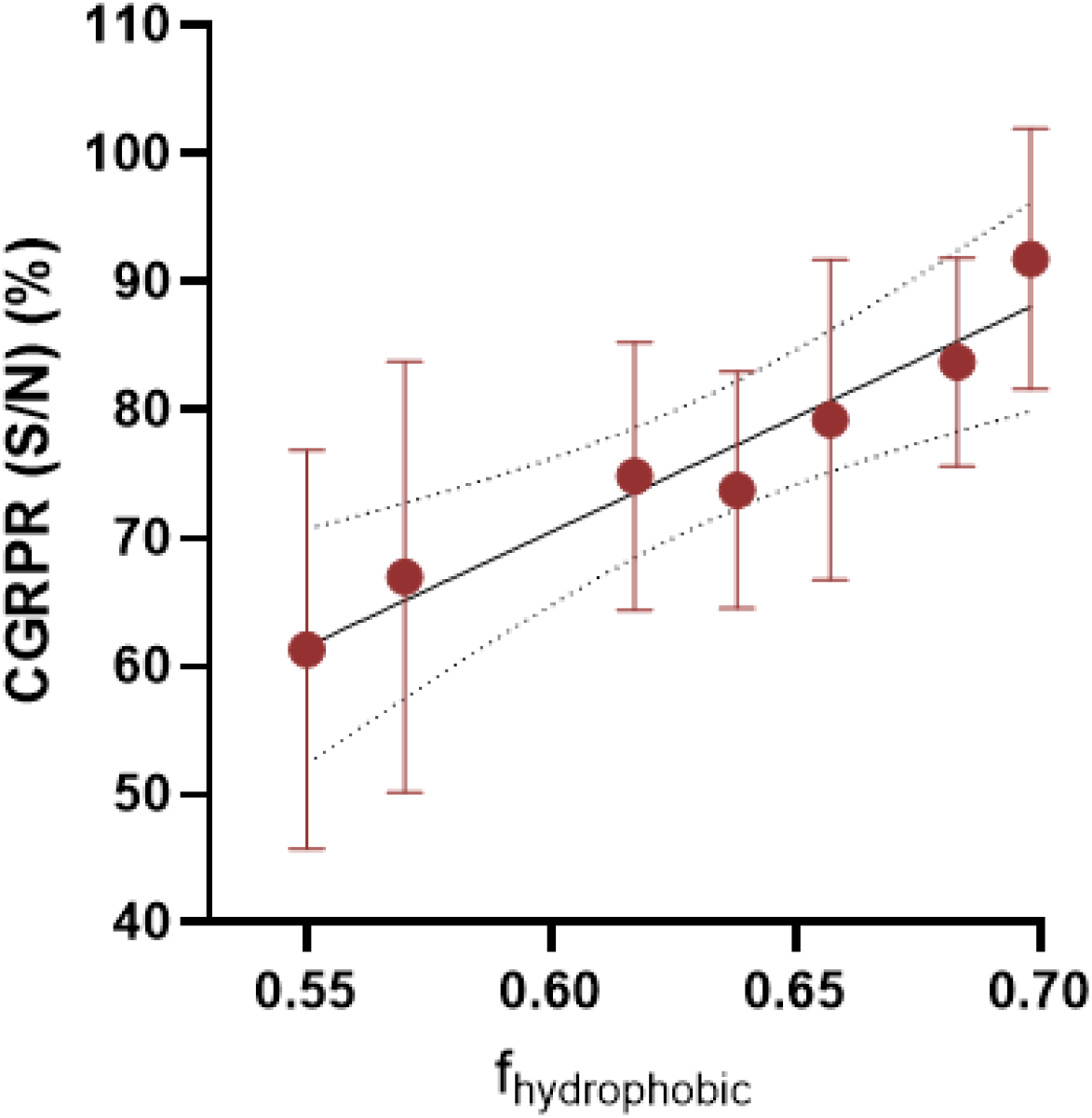
Correlation between the CGRPR solubilisation efficiency and the hydrophobicity of the BzAM series (BzAM10 – BzAM40). Solubilisation efficiency is the percentage CGRPR in the original membrane preparation detected in the soluble fraction (S/N), determined by densitometric analysis of western blots. Data are mean ± s.e.m. (n = 3). The total hydrophobic molar fraction (f_hydrophobic_) of each terpolymer consists of f_BZ_ (N-benzyl maleimide fraction) and f_St_ (styrene fraction). The broken curved lines show the 95% confidence limits of the fitted straight line.

### 3.2. Optimising Amphiphilic Polymer Properties

Efficient membrane solubilisation by amphiphilic polymers necessitates a precise balance between their hydrophilic and hydrophobic characteristics to facilitate interaction at the interface between the aqueous phase and the hydrophobic lipid bilayer. Amphiphilic polymers must have sufficient hydrophilicity to maintain aqueous solubility while exhibiting adequate hydrophobicity to enable bilayer insertion, membrane disruption, and subsequent nanodisc formation. Membrane solubilisation occurs through the insertion of the polymer’s hydrophobic moieties into the lipid bilayer, disrupting the hydrophobic interactions between lipid acyl chains and leading to the extraction of membrane segments into nanoscale discs. Following nanodisc formation, the polymer encompasses the disc periphery, with its hydrophobic segments intercalating between lipid acyl chains (14,35).

Achieving an optimal balance between hydrophobicity and hydrophilicity is critical. Excessive hydrophobicity promotes polymer aggregation and precipitation, whereas excessive hydrophilicity compromises membrane insertion and solubilisation efficiency. This balance is exemplified by the limitations of SMA1:1, which, due to its insufficient hydrophobicity, exhibits poor bilayer interaction and low solubilisation efficiency. Conversely, highly hydrophobic SMA variants (*e*.*g*., SMA4:1) lack sufficient hydrophilicity to establish the initial electrostatic interactions with lipid headgroups necessary for membrane association (36). The stepwise hydrophobic adjustments in the BzAM terpolymer series illustrate the important role of the amphiphilic balance in optimising membrane protein solubilisation.

### 3.3 CGRP Receptor Ligand Binding Capability Is Preserved In BzAM Nanodiscs

Efficient extraction of the target membrane protein from its native lipid bilayer environment is evidently a prerequisite for any candidate solubilising polymer; however, this is not the only criterion that must be met. Having established that BzAM terpolymers were effective at solubilising the CGRPR, it was important to confirm that the extracted receptor retained its functional ligand-binding ability when encapsulated as CGRPR-BzAM lipid particle (BzAMLP). This was assessed using NanoBRET proximity assay in which the binding of a fluorescent ligand (BODIPY-CGRP) to the NLuc-CGRPR receptor was monitored by BRET (24,37). NLuc-tagged GPCRs and BODIPY630/650-tagged fluorescent ligands form an established and effective BRET donor-acceptor pair for NanoBRET ligand binding assays (38).

HEK 293S GnTI^−^ TetR cells were engineered to stably overexpress inducible NLuc-CGRPR. Using BzAM35 as a representative polymer from the BzAM series, NLuc-CGRPR was solubilised and the NLuc-CGRPR-BzAMLP nanodiscs generated were purified from the soluble fraction using immobilised metal ion affinity chromatography (affinity purification). Samples obtained at each stage of the purification process, analysed by western blotting (for NLuc-CGRPR) (Figure 5A) and SDS-PAGE (for total protein) (Figure 5B), confirmed enrichment of NLuc-CGRPR.

**Figure 5.**
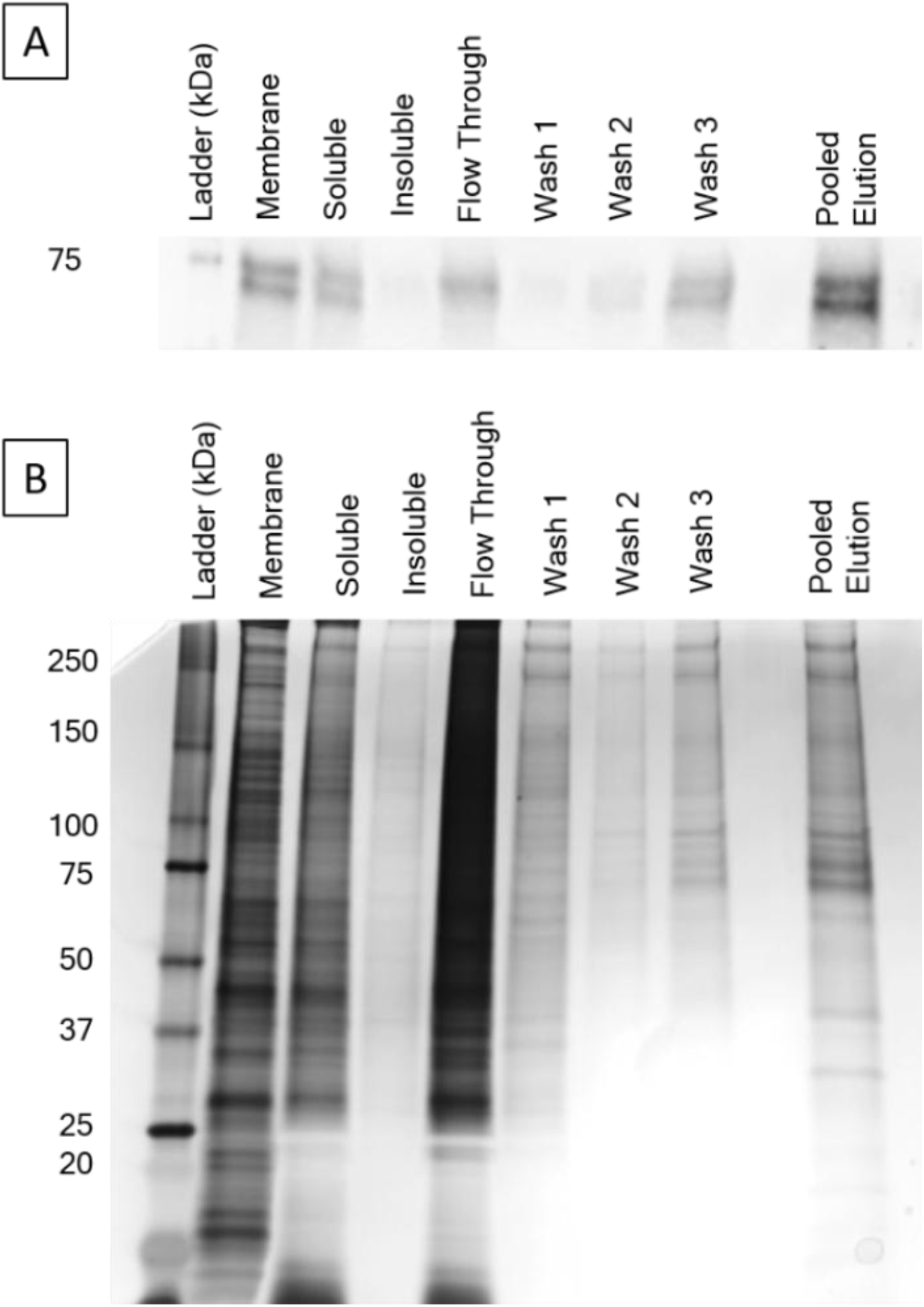
Enrichment of Nluc-CGRPR-BzAMLPs using Ni^2+^-NTA. CGRPR was solubilised using 2.5% (w/v) BzAM35. The CGRPR-BzAMLPs were enriched using Ni^2+^-NTA. (A) Western blot detection (anti-HA antibody) of CGRPR in fractions from the enrichment of Nluc-CGRP-BzAMLP using Ni^2+^-NTA. (B) Silver stain of fractions from the enrichment of Nluc-CGRP-BzAMLP using Ni^2+^-NTA. Data are representative of three independent experiments.

The Nluc-CGRPR-BzAMLPs were assessed for the retention of high-affinity ligand binding capability using the fluorescent CGRP analogue BODIPY-CGRP in a BRET-based assay. Saturation ligand binding with increasing concentrations of BODIPY-CGRP (Figure 6) established retention of high-affinity ligand binding by the NLuc-CGRPR when encapsulated in the BzAMLP nanodiscs, with an affinity (pK_d_ = 9.6 ± 0.009, mean ± s.e.m., n = 3) comparable to published values for CGRP binding to the CGRPR (IUPHAR/BPS Guide to Pharmacology, calcitonin receptors; www.guidetopharmacology.org). The demonstration that the BzAM terpolymer solubilises the CGRPR, a dynamic heterodimeric GPCR, with retention of high-affinity binding, will open new opportunities for research exploiting polymers to study membrane proteins encapsulated in nanodiscs.

**Figure 6.**
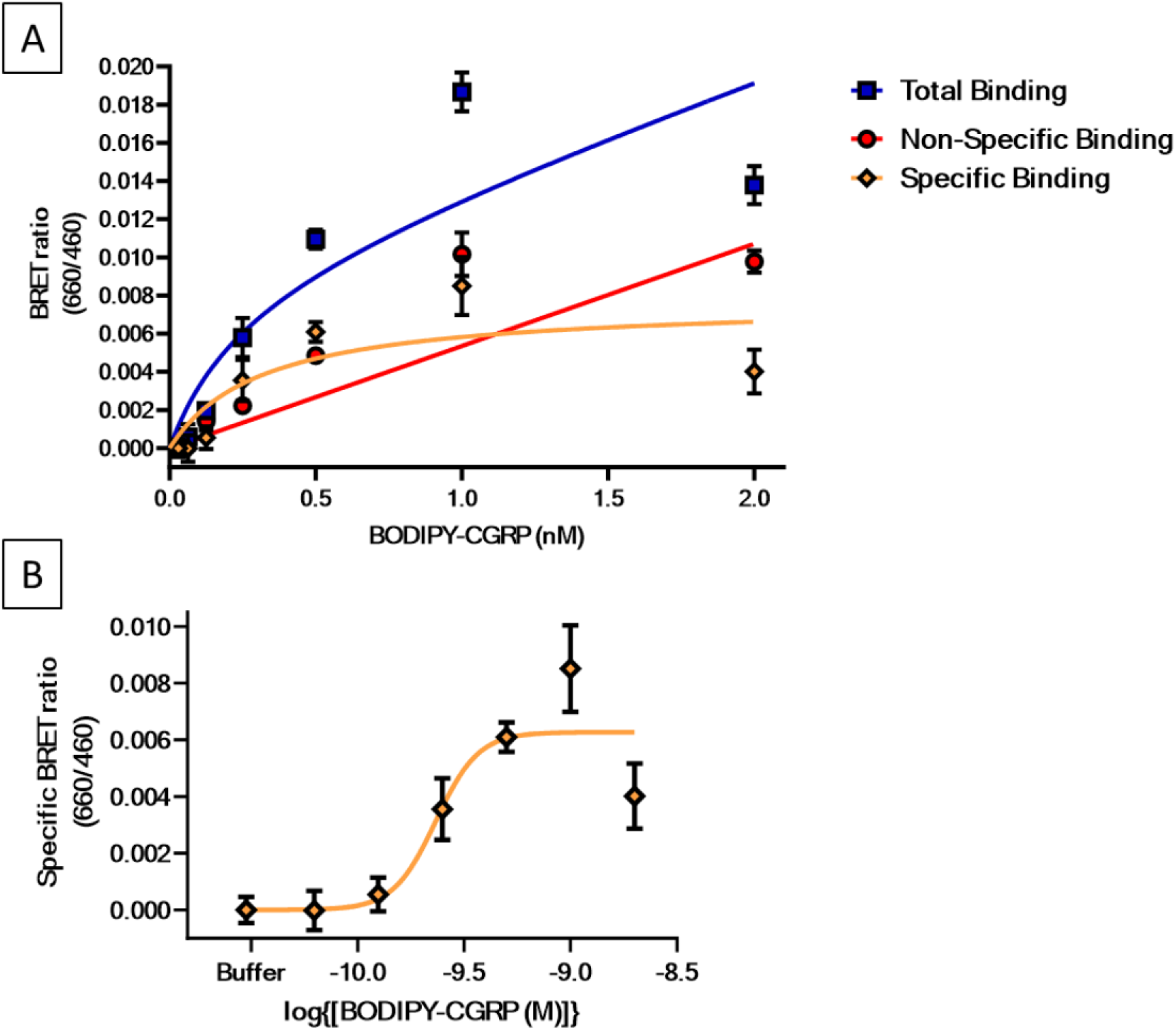
Ligand binding capability of CGRPR preserved in a BzAMLP. (A)Total binding, non-specific binding and specific binding of BODIPY-CGRP to Nluc-CGRPR-BzAMLPs. Non-specific binding was defined by olcegepant (10 nM). (B)Specific binding of BODIPY-CGRP to Nluc-CGRPR-BzAMLPs. Data are mean ± s.e.m. (n = 3).

For any agent used to solubilise a membrane protein of interest, it is important to determine the effect that agent has on the function of the membrane protein, as the solubilisation procedure can markedly alter protein behaviour. A good example of this was demonstrated for the muscarinic receptor family of GPCRs. Even within membranes, experimental conditions such as buffer composition or temperature can markedly change antagonist selectivity (39). These changes become even more marked after solubilisation with detergent. For example, muscarinic receptors isolated from rat myocardium, which are now known to belong to the M2 subtype, have an M1-subtype pharmacology after solubilisation with digitonin (40). Another pattern of pharmacology is seen after solubilisation with the zwitterionic detergent 3-(3-cholamidopropyl)dimethylammonio-2-hydroxy-1-propane sulfonate (CHAPSO). Interestingly, it proved possible to swap between the digitonin and CHAPSO pharmacology simply by exchanging detergents by gel chromatography (41). These studies show that the immediate environment in which a membrane protein resides can have important implications for its functional characteristics; it simply cannot be assumed that solubilising a receptor will give native function. Whilst the above studies were carried out with detergents, similar effects are likely to be at work for polymers, as these will influence the physical properties of the encapsulated lipid bilayer, such as fluidity, lipid packing and lateral pressure. Indeed, we and others have reported polymer-specific effects on the encapsulated protein in nanodiscs. For example, light-induced changes in rhodopsin conformation were dictated by the specific polymer employed (42), likewise for the photoreceptor/transducer complex (NpSRII2/NpHtrII2) from *Natronomonas pharaonis* (43). Consequently, it is important to have a wide range of polymers with different architectures available to facilitate optimal solubilisation of the protein of interest while retaining its functional capability. The BzAM series therefore, provides a novel resource for studying membrane proteins in nanodiscs.

## 4. CONCLUSION

This study establishes the utility of BzAM terpolymers for detergent-free solubilisation and functional analysis of GPCRs. These terpolymers effectively extracted the human CGRP receptor from mammalian membranes while preserving its high-affinity ligand-binding capability, exhibiting a positive correlation between polymer hydrophobicity and membrane solubilisation efficiency. As the diversity of membrane-active polymers continues to expand [reviewed in (44)], it becomes increasingly important that such materials not only enable efficient extraction and retention of protein function but are also tailored for downstream applications and biophysical analyses. These additional requirements will guide the targeted design of polymers tailored to distinct applications. This will include the development of novel polymers in which a specific aspect of the polymer is systematically altered to provide finely-tuned variation in a single parameter. The BzAM terpolymer series exemplifies this principle, providing a series of well-defined, tuneable polymers in which a single molecular parameter, in this case hydrophobicity, is systematically varied in a step-wise manner. This provides a level of subtle versatility not previously available for studying membrane proteins in near-native nanodiscs. Insights from this work will inform the development of polymer-based strategies optimised for structural and mechanistic studies, including investigations of lipid-modulated GPCR conformational dynamics, and future high-throughput drug discovery applications.

## Acknowledgements

The authors acknowledge funding from the Wellcome Trust via a Technology Development grant entitled “Unshackling Membrane Protein Research: New Amphiphilic Copolymers for Extraction of Stable, Active Membrane Proteins” (Grant No. 223728/Z/21/Z).

## COI/Notes

The authors declare the following competing financial interest(s): GCK and BK are affiliates of Nanosene (Pty) Ltd, a company that commercialises amphiphilic polymers for membrane protein research and beyond.

## Abbreviations

BV: bed volume
BODIPY: boron-dipyrromethene
BRET: bioluminescence resonance energy transfer
BzAM: poly(styrene-*co*-maleic acid-*co*-(*N*-benzyl)maleimide)
BzAMLP: poly(styrene-*co*-maleic acid-*co*-(*N*-benzyl)maleimide) lipid particle
cDNA: complementary DNA
CGRP: calcitonin gene-related peptide
CGRPR: calcitonin gene-related peptide receptor
CHAPSO: 3-(3-cholamidopropyl)dimethylammonio-2-hydroxy-1-propane sulfonate
CLR: calcitonin receptor-like receptor
DMEM/F12: Dulbecco’s Modified Eagle Medium/Nutrient Mixture F-12 FBS foetal bovine serum
FRET: Förster Resonance Energy Transfer
G418: geneticin
GNTI: N-acetylglucosaminyltransferase I deficient
GPCRs: G-protein-coupled receptors
HA: hemagglutinin
HEK 293S: human embryonic kidney 293S cells
HEPES: 4-(2-hydroxyethyl)-1-piperazineethanesulfonic acid His_10_ deca-histidine
NanoBRET: Nanoluciferase Bioluminescence Resonance Energy Transfer Ni-NTA nickel nitrilotriacetic acid
NLuc: Nanoluciferase
P2A: porcine teschovirus-1 2A element
Pra: propargylglycine
RAFT: reversible addition−fragmentation chain transfer
RAMP1: receptor activity modifying protein 1
SDS: sodium dodecyl sulfate
SDS: PAGE SDS polyacrylamide gel electrophoresis SMA poly(styrene-*co*-maleic acid)
SMAnh: poly(styrene-*alt*-maleic anhydride)
TetR: tetracycline repressor

